# Cochlear Neurotrophin-3 overexpression at mid-life prevents age-related cochlear synaptopathy and slows age-related hearing loss

**DOI:** 10.1101/2022.04.05.487225

**Authors:** Luis R. Cassinotti, Lingchao Ji, Beatriz C. Borges, Nathan D. Cass, Aditi S. Desai, David C. Kohrman, M. Charles Liberman, Gabriel Corfas

**Author notes:** Correspondence: Gabriel Corfas, PhD, Medical Sciences I Building, Rm. 5428, 1150 West Medical Center Drive, Ann Arbor, MI 48109-5616. Peking University Shenzhen Hospital, Futian District, Shenzhen, Guangdong Province, China, 518036 (Lingchao Ji). The Otology Group of Vanderbilt; Vanderbilt University Medical Center Department of Otolaryngology; Nashville, TN.

## Abstract

Age-related hearing loss (ARHL) is the most prevalent sensory deficit in the elderly. This progressive pathology often has psychological and medical comorbidities, including social isolation, depression, and cognitive decline. Despite ARHL’s enormous impact, no therapies to prevent or slow its progression exist. Loss of synapses between inner hair cells (IHCs) and spiral ganglion neurons (SGNs), a.k.a. cochlear synaptopathy, is an early event in ARHL, preceding neuronal and hair cell loss. To determine if age-related cochlear synaptopathy can be prevented, and if this impacts the time-course of ARHL, we tested the effects of cochlear overexpression of neurotrophin-3 (Ntf3) starting at middle-age. We chose Ntf3 because this neurotrophin regulates the formation of IHC-SGN synapses in the neonatal period. We now show that triggering Ntf3 overexpression by cochlear supporting cells starting in middle age rapidly increases the amplitude of sound-evoked neural potentials, indicating that Ntf3 improves cochlear function when pathology is minimal. Furthermore, near the end of their lifespan, Ntf3-overexpressing mice have milder ARHL, with enhanced cochlear neural function and reduced cochlear synaptopathy. Our results provide evidence that age-related decrease in cochlear Ntf3 expression contributes to ARHL. and Ntf3 supplementation could serve as a therapeutic for this prevalent disorder. Furthermore, these findings suggest that increased expression of factors that regulate synaptogenesis during development might prevent age-related synaptopathy, a process involved in several central nervous system degenerative disorders.

## Introduction

Age-related hearing loss (ARHL), also known as presbycusis, is the most prevalent sensory deficit in the elderly, affecting a third of people over age 65 and half of those over 85 in the US (Agrawal, Platz, & Niparko, 2008; Bainbridge & Wallhagen, 2014). This progressive pathology is characterized by elevated hearing thresholds and difficulties with speech comprehensions especially in noisy environments. It is often associated with psychological and medical comorbidities, including social isolation, frailty, depression, and cognitive decline (Livingston et al., 2020; Powell, Oh, Lin, & Deal, 2021). Despite ARHL’s enormous societal and economic impact (Huddle et al., 2017), no therapies to prevent or slow this process exist.

While the cellular and molecular mechanisms of ARHL remain unclear, studies in mice and humans demonstrated that loss of synapses between inner hair cells (IHCs) and spiral ganglion neurons (SGNs), a.k.a. cochlear synaptopathy, is an early event in ARHL, preceding hair cell (HC) loss (Makary, Shin, Kujawa, Liberman, & Merchant, 2011; Sergeyenko, Lall, Liberman, & Kujawa, 2013; Viana et al., 2015; Wu et al., 2019). Furthermore, studies in mice showed that noise-induced cochlear synaptopathy at an early age accelerates the onset of SGN loss and age-related threshold increases (Kujawa & Liberman, 2006). Based on these observations, we wondered if cochlear synaptopathy can be prevented or delayed, and if achieving that alters the progression of ARHL. For this, we focused on neurotrophin-3 (Ntf3), a neurotrophic factor critical for the survival of SGNs during embryogenesis (Ernfors, Van De Water, Loring, & Jaenisch, 1995; Farinas, Jones, Backus, Wang, & Reichardt, 1994) which we showed regulates the formation of IHC-SGN synapses in the neonatal period and induces their regeneration and functional recovery after acoustic injury in young mice (Wan, Gomez-Casati, Gigliello, Liberman, & Corfas, 2014). Moreover, using a reporter mouse in which β-galactosidase is co-expressed with Ntf3 (Farinas et al., 1994), we showed that cochlear Ntf3 expression decreases during aging (Sugawara, Murtie, Stankovic, Liberman, & Corfas, 2007), suggesting that lower Ntf3 levels could contribute to ARHL.

Here we tested the impact of Ntf3 overexpression on ARHL in mice using transgenic Ntf3 overexpression starting at middle age. We show that induction of IHC supporting-cell Ntf3 overexpression in mice at 1 year of age rapidly increases the amplitude of sound evoked potentials, indicating that Ntf3 treatment can improve cochlear function in mice with mild hearing impairment. Furthermore, Ntf3 overexpression slows the progression of ARHL, resulting in enhanced cochlear function and reduced age-related cochlear synaptopathy in mice approaching the end of their lifespan.

## Results

To determine the impact of increasing supporting cell Ntf3 expression on the time-course of ARHL, we studied two cohorts of mice. One group (*Ntf3*^*Stop*^*:Slc1a3/CreER*^*T*^) carried an inducible Ntf3 transgene (*Ntf3*^*Stop*^) (Wan et al., 2014) and the *CreER*^*T*^ recombinase transgene under the control of the GLAST (*Slc1a3*) promoter (Y. Wang et al., 2012) to drive Ntf3 overexpression in inner border and inner phalangeal cells, the supporting cells immediately surrounding the IHCs and their synaptic connections with auditory nerve fibers. The control group consisted of littermates carrying only the *Ntf3*^*Stop*^ transgene (Wan et al., 2014). At 59 weeks of age, we measured baseline auditory function in both cohorts by recording distortion product otoacoustic emissions (DPOAEs) and auditory brainstem responses (ABRs) (Fig. 1A). DPOAEs reflect outer hair cell (OHC) function; ABRs reflect the function of the ascending auditory system, from the activation of IHCs and SGNs to the inferior colliculus (Kohrman, Borges, Cassinotti, Ji, & Corfas, 2021). The groups exhibited no differences in DPOAE nor in ABR thresholds at the baseline time point (Fig 1B and C). Furthermore, at this time point, the groups did not differ in the amplitudes of the first peak of the ABR waveform (Figs. 1D and 2A), which reflects the synchronous activity of SGN fibers in response to sound. Analysis of the later peaks in the ABR waveforms (Fig. 2B) indicated that the signaling along the ascending auditory pathway at baseline was similar between the groups as well.

**Figure 1:**
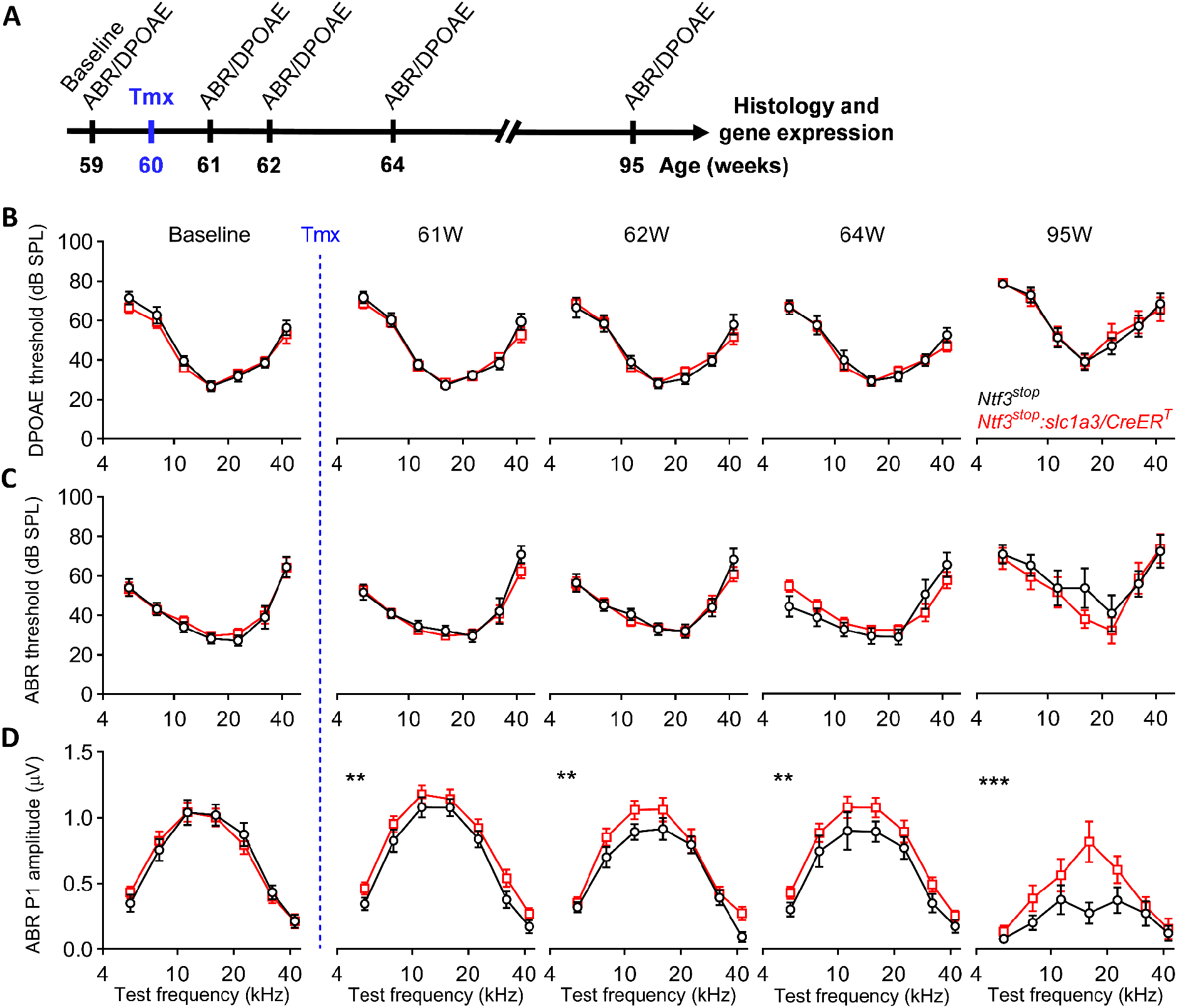
Ntf3 overexpression at mid-life slows the age-related decline in ABR peak 1 amplitude but does not alter ABR and DPOAE shifts. **(A)** Timeline of the experimental design. Ntf3 overexpressing (*Ntf3*^*stop*^*:Slc1a3/CreER*^*T*^) and control (*Ntf3*^*stop*^) mice were used in this study. Ntf3 overexpression was induced by intraperitoneal tamoxifen injection at 60 weeks of age (highlighted in blue). Baseline ABRs and DPOAEs were recorded one week before tamoxifen treatment (59 weeks of age) and at four later time points (61, 62, 64 and 95 weeks of age). After the last ABR and DPOAE tests, cochleae were collected and processed for either immunohistochemistry followed by confocal microscopy to evaluate HCs and IHC synapse counts or for RT-qPCR to evaluate Ntf3 mRNA expression. **(B)** Ntf3 overexpression starting at 60 weeks of age does not alter DPOAE thresholds during the following 35 weeks. **(C)** Ntf3 overexpression starting at 60 weeks of age does not alter ABR thresholds during the following 35 weeks. **(D)** Ntf3 overexpression starting at 60 weeks of age increases ABR peak 1 amplitudes one week later and slows the decline of ABR peak 1 amplitudes at 80 dB SPL during the following 34 weeks. *Ntf3*^*stop*^ n = 7-13; *Ntf3*^*stop*^*:Slc1a3/CreER*^*T*^ n = 7-17. * p < 0.05; ** p < 0.01; *** p < 0.001 by two-way ANOVA, followed by Bonferroni’s multiple comparisons test. Error bars represent SEM.

**Figure 2.**
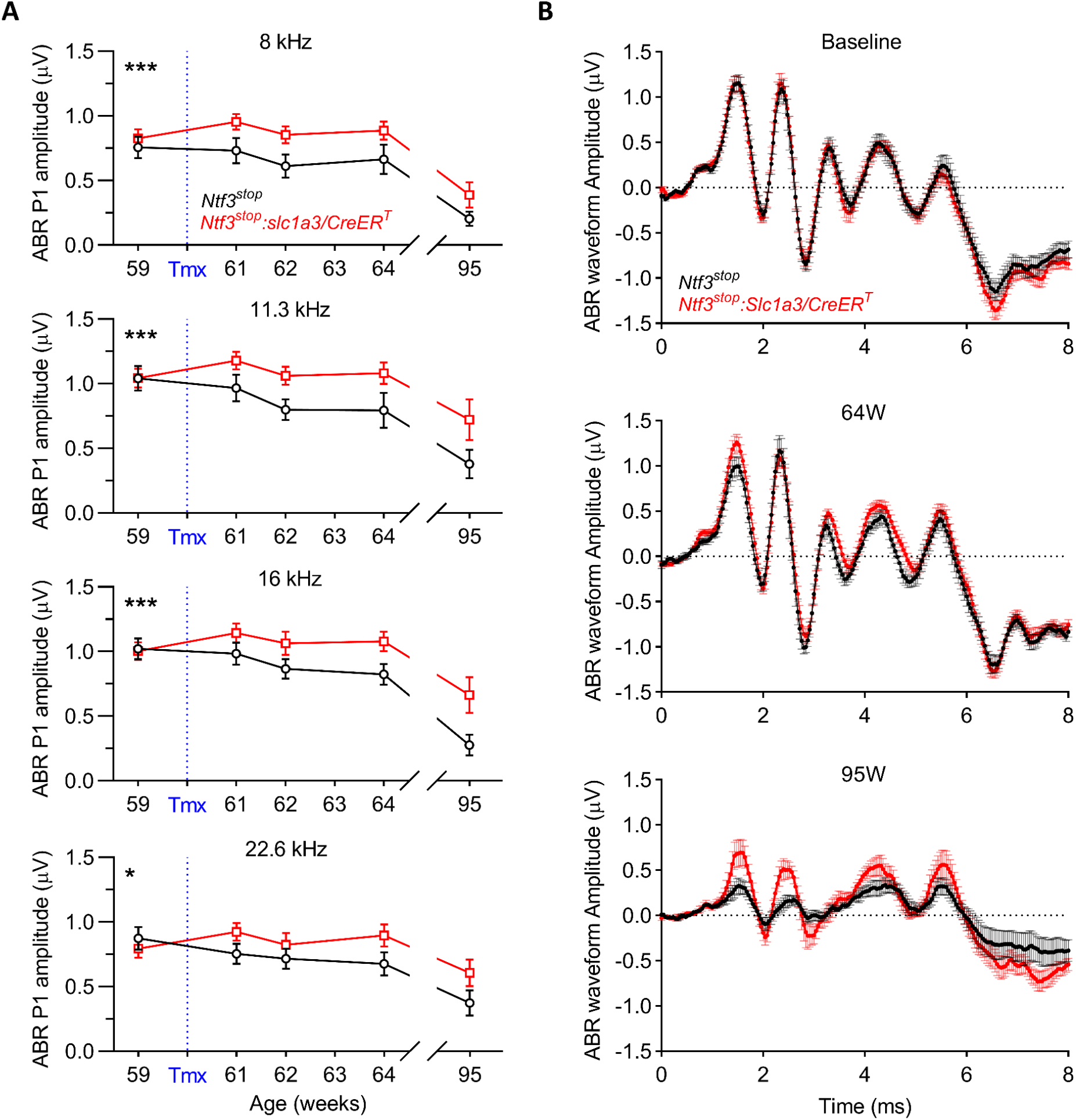
Ntf3 overexpression improves sound-evoked activity along the cochlear axis and in higher auditory centers in the aging brain. **(A)** The effects of Ntf3 overexpression starting at 60 weeks of age is evident along the cochlear axis. **(B)** Mean ABR waveforms recorded at baseline, 64 and 95 weeks of age show that Ntf3 overexpression increases sound-induced signaling along the ascending auditory pathway in aging mice. At baseline, before tamoxifen treatment, no intergroup differences were seen in ABR amplitudes. At 64 weeks, peak 1 amplitudes were increased in Ntf3 overexpressing mice. All ABR peaks were reduced in both cohorts by 95 weeks, but Ntf3 overexpressors exhibited smaller decreases at this age. ABRs shown here are group means in response to 16 kHz tone pips at 80 dB SPL. *Ntf3*^*stop*^ n = 7-13; *Ntf3*^*stop*^*:Slc1a3/CreER*^*T*^ n = 7-17. * p < 0.05; *** p < 0.001 by two-way ANOVA. Error bars represent SEM.

All animals were treated with tamoxifen at 60 weeks of age to induce *CreER*^*T*^-mediated activation of the *Ntf3*^*Stop*^ transgene, which occurs only in mice carrying the *Slc1a3/CreER*^*T*^ allele, and auditory function was followed longitudinally in both cohorts. Notably, one week after activation of Ntf3 overexpression (61 weeks of age), the suprathreshold ABR peak 1 amplitudes in *Ntf3*^*Stop*^*:Slc1a3/CreER*^*T*^ mice were higher than in the control group (Fig. 1D, 2A and 3A), indicating that increased Ntf3 availability leads to an acute increase in the sound-evoked potentials mediated by IHC-SGN synapses. This contrasts with what we found in young (8-week-old) mice, where activation of the Ntf3 transgene does not alter ABRs or DPOAEs (Fig. 3B), indicating that the acute effects of Ntf3 are age dependent.

**Figure 3.**
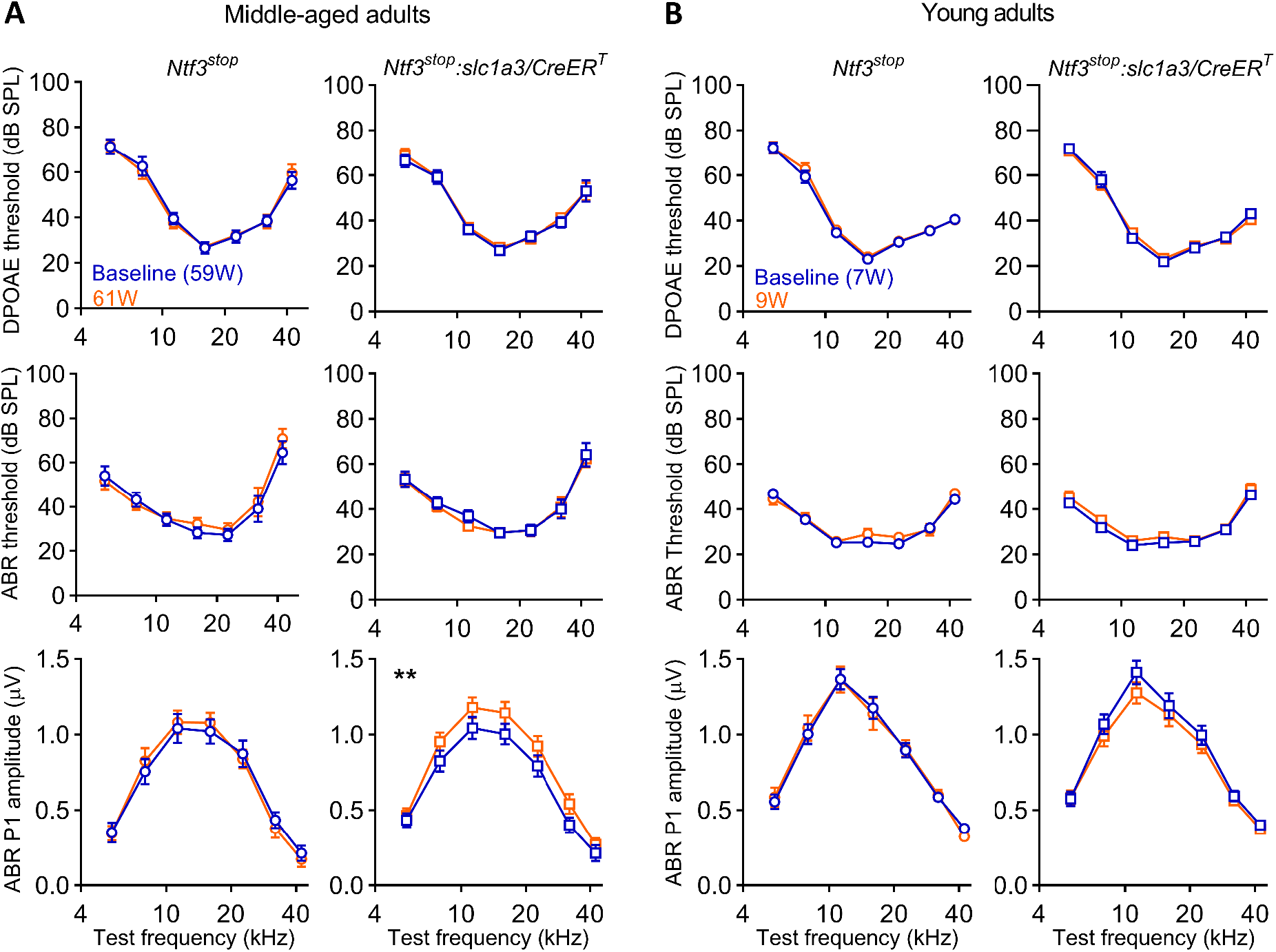
Ntf3 overexpression acutely increases ABR peak 1 amplitude in middle-aged mice but not in the young. **(A)** DPOAE and ABR recordings, before and one-week after tamoxifen injection in middle-aged (60-week-old) show that tamoxifen treatment does not alter ABR and DPOAE thresholds but acutely increases ABR peak 1 amplitudes in the Ntf3 overexpressing mice. **(B)** No increase was seen when young adult (8-week-old) were tested. Suprathreshold ABR peak 1 amplitudes were analyzed at 80 dB SPL. Middle-aged adults *Ntf3*^*stop*^ n = 11; Middle-aged *Ntf3*^*stop*^*:Slc1a3/CreER*^*T*^ n = 13-17; Young adults *Ntf3*^*stop*^ n = 22; Young adults *Ntf3*^*stop*^*:Slc1a3/CreER*^*T*^ n = 20. * p < 0.05; ** p < 0.01 by two-way ANOVA, followed by Bonferroni’s multiple comparisons test. Error bars represent SEM. and young adult mice.

As the animals aged, both groups showed similar increases in DPOAE and ABR thresholds (Fig. 1B and 1C). However, suprathreshold ABR peak 1 amplitudes were higher in *Ntf3*^*Stop*^*:Slc1a3/CreER*^*T*^ than in the *Ntf3*^*Stop*^ controls at each post-tamoxifen time point (Figs 1D) and in every region of the cochlea (2A and 2B), indicating that Ntf3 overexpression preserves the connections between IHCs and SGNs during aging along the whole cochlear axis. Furthermore, Ntf3 overexpression also enhanced the later peaks in the ABR waveforms at 95 weeks of age (Fig. 2B), showing that the Ntf3-mediated improvement in cochlear function is also reflected in higher synchronous activation along the ascending auditory pathway in the aging brain.

After the last session of auditory testing, inner ears were harvested and used for molecular and histological analysis (Fig. 1A). As expected from our prior studies on young adult mice (Wan et al., 2014), Ntf3 mRNA levels were significantly higher in the 95-week-old *Ntf3*^*Stop*^*:Slc1a3/CreER*^*T*^ mice compared to their 95-week-old littermates carrying only the *Ntf3*^*Stop*^ transgene (Fig. 4). Furthermore, consistent with our prior studies (Sugawara et al., 2007), Ntf3 levels in control mice at 95 weeks were ∼40% lower than those in 10-week-old controls. Notably, Ntf3 levels in *Ntf3*^*Stop*^*:Slc1a3/CreER*^*T*^ mice were not statistically different from those seen in young *Ntf3*^*Stop*^ mice, indicating that tamoxifen induction was sufficient to maintain Ntf3 expression in the old *Ntf3*^*Stop*^*:Slc1a3/CreER*^*T*^ mice at the same level as they had in their youth.

**Figure 4.**
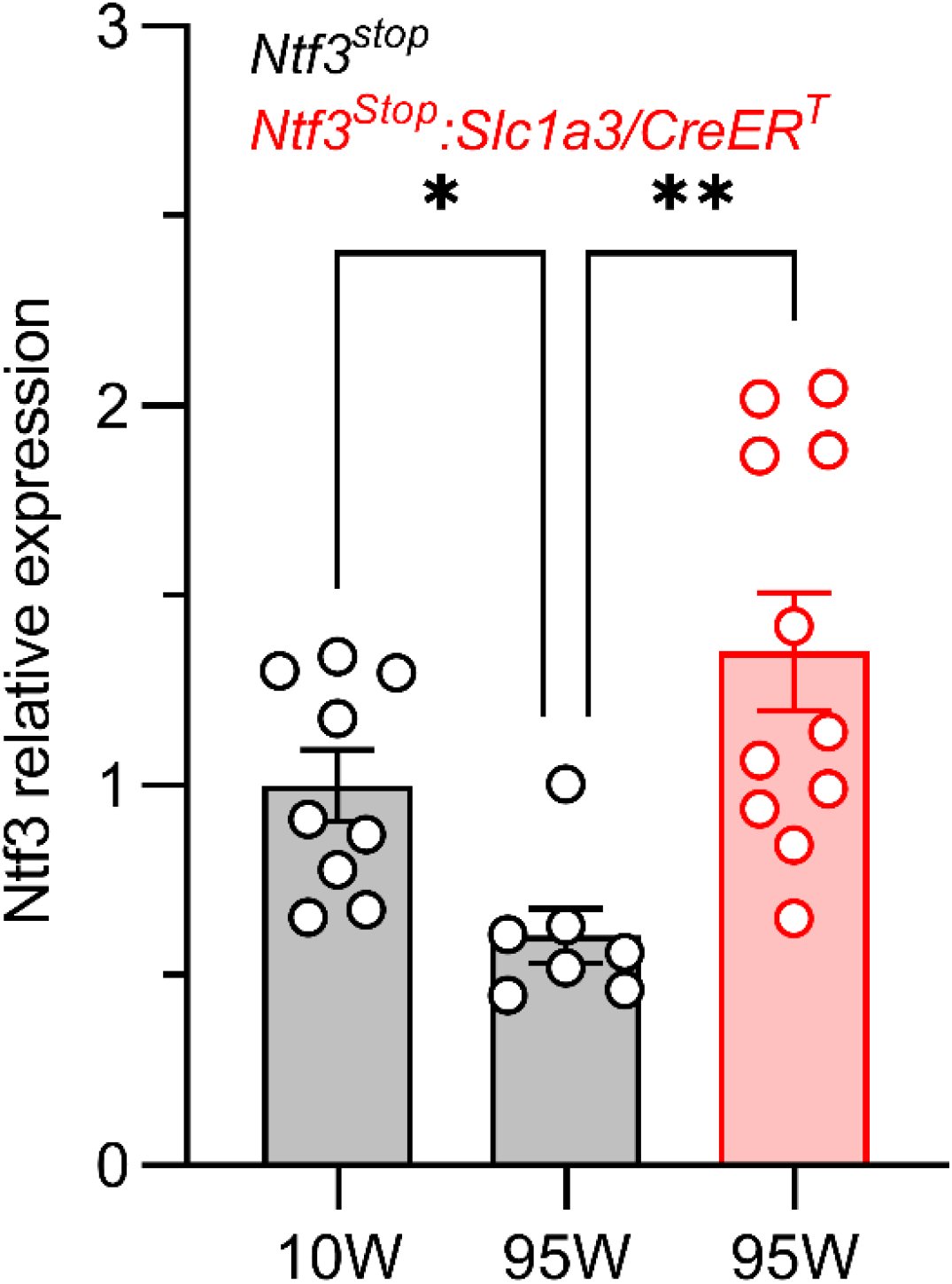
Cochlear Ntf3 mRNA levels decrease by 40% between 10 and 95 weeks of age in control mice, but expression levels in 95-week-old *Ntf3*^*stop*^*:Slc1a3/CreER*^*T*^ mice reach those seen in young mice when tamoxifen is injected at 60 weeks of age. RT-qPCR evaluation of Ntf3 mRNA levels in cochleae from control (Ntf3^stop^) mice at 10 and 95 weeks of age (grey bars) and in Ntf3^stop^:Slc1a3/CreER^T^ mice at 95 weeks of age having treated with tamoxifen at 60 weeks. 10-week-old *Ntf3*^*stop*^ n = 9; 95-week-old *Ntf3*^*stop*^ n = 7; 95-week-old *Ntf3*^*stop*^*:Slc1a3/CreER*^*T*^ n = 11. * p < 0.05; ** p < 0.01 by Kruskal-Wallis test, followed by Dunn’s multiple comparisons test. Error bars represent SEM.

Congruent with the preservation of peak 1 amplitudes in 95-week-old *Ntf3*^*Stop*^*:Slc1a3/CreER*^*T*^ mice, their IHC-SGN synapse counts were also comparable to those found in 10-week-old *Ntf3*^*Stop*^ mice, and significantly higher than in the 95-week-old *Ntf3*^*Stop*^ littermates (Fig. 5A and 5B), demonstrating that Ntf3 overexpression was able to prevent age-related synaptopathy. Furthermore, we found a significant correlation between Ntf3 mRNA levels, synapse counts and ABR peak 1 amplitudes at 95 weeks of age (Fig. 5C; ABR peak 1 amplitudes vs. Ntf3 expression level r = 0.846 p < 0.001; IHC synapses vs. Ntf3 expression level r = 0.733 p = 0.0311 and IHC synapses vs. ABR peak 1 amplitudes r = 0.783 p = 0.017). In contrast, in agreement with the DPOAE thresholds shifts seen in both genotypes at 95 weeks (Fig. 1B) and the extent of OHC survival previously observed at this age (Sergeyenko et al., 2013), the OHC survival was similar in *Ntf3*^*Stop*^*:Slc1a3/CreER*^*T*^ and *Ntf3*^*Stop*^ mice (Fig. 6), indicating that supporting cell Ntf3 overexpression protects IHC-SGN synapses but not OHCs, which do not express the receptor for Ntf3 (TrkC) and only transiently express the receptor for Bdnf (trkB) during the neonatal period (Knipper, Zimmermann, Rohbock, Kopschall, & Zenner, 1996; Ylikoski et al., 1993).

**Figure 5.**
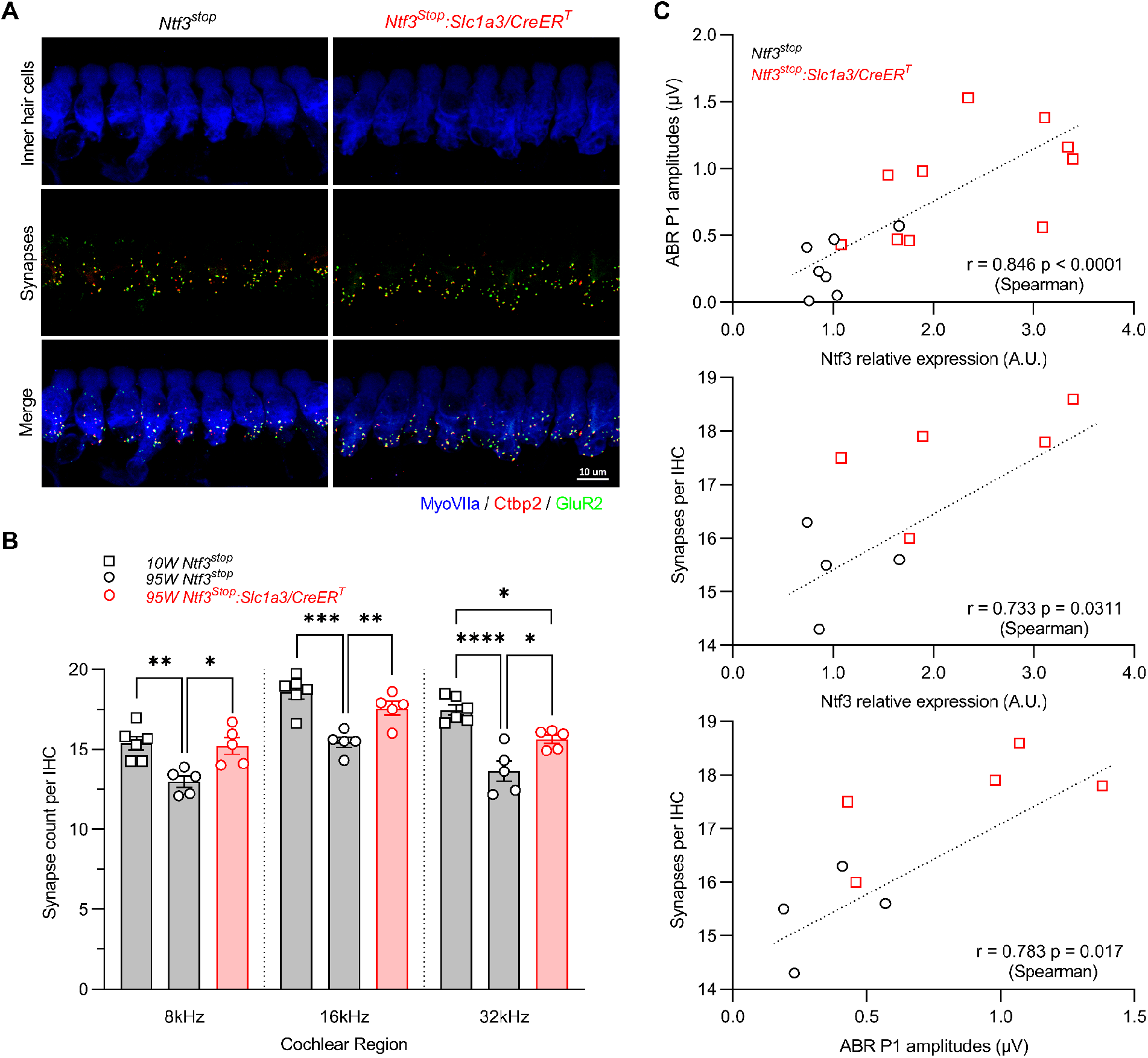
Increased Ntf3 expression levels at 95 weeks of age correlates with increased IHC synapse and ABR peak 1 amplitude. **(A)** Representative confocal images of IHC synapses at the 16 kHz region of a 95-week-old *Ntf3*^*stop*^ and *Ntf3*^*stop*^*:Slc1a3/CreER*^*T*^ cochlea immunolabeled for hair cells (MyoVIIa), pre-synaptic ribbons (Ctbp2) and post-synaptic receptor patches (GluR2). **(B)** At 95 weeks of age, Ntf3 overexpressors have 14-17% more IHC synapses compared to controls, reaching synapse densities comparable to those seen in 10-week-old controls. 10-week-old *Ntf3*^*stop*^ n = 6; 95-week-old *Ntf3*^*stop*^ n = 5; 95-week-old *Ntf3*^*stop*^*:Slc1a3/CreER*^*T*^ n = 5. * p < 0.05; ** p < 0.01; *** p < 0.001; **** p < 0.0001 by one-way ANOVA. Error bars represent SEM. **(C)** Ntf3 expression levels, ABR peak 1 amplitudes, and IHC synapses density strongly correlate in 95-week-old mice. Correlations were evaluated using non-parametric Spearman correlation test. p < 0.05 were considered as statistically significant differences.

**Figure 6.**
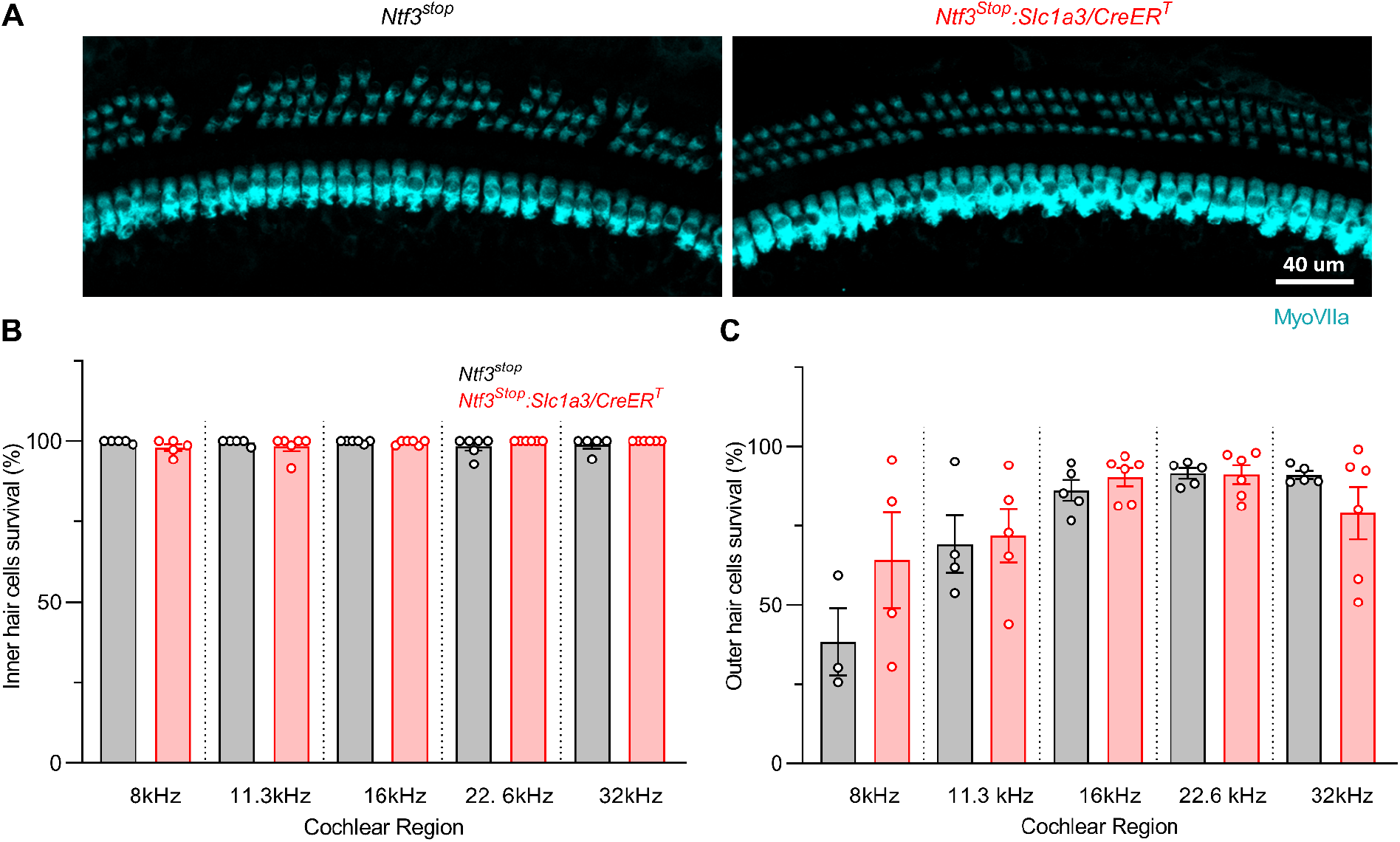
Ntf3 overexpression starting at mid-life does not alter age-related OHC loss. **(A)** Representative confocal images of the 16 kHz region of *Ntf3*^*stop*^ and *Ntf3*^*stop*^*:Slc1a3/CreER*^*T*^ cochleae immunolabeled for hair cells (MyoVIIa) showing mild OHC loss without IHC loss at 95 weeks of age. **(B)** Quantitative analysis shows no IHC loss. **(C)** Both groups show equal degrees of mild OHC loss. *Ntf3*^*stop*^ n = 3-6; *Ntf3*^*stop*^*:Slc1a3/CreER*^*T*^ n = 5-6. Non-significant differences were found by two-tailed t-test within the evaluated cochlear frequencies. Error bars represent SEM.

## Discussion

ARHL has reached epidemic proportions (Agrawal et al., 2008; Bainbridge & Wallhagen, 2014), and hearing impairment has been shown to contribute to cognitive decline and dementia (Livingston et al., 2020; Powell et al., 2021). Thus, developing therapies to prevent ARHL is an important public health priority. Our results provide the first demonstration that increasing Ntf3 levels in the middle-aged cochlea promotes the maintenance of IHC-SGN synapses and their function during murine aging, supporting the potential of Ntf3 supplementation as a powerful strategy to reduce the burden of ARHL in humans. However, the increased Ntf3 expression and IHC synapse preservation does not prevent OHC loss and the consequent age-related cochlear threshold shifts. These findings indicate that OHC survival during aging does not depend on Ntf3, SGNs, or IHC-SGN synapses, and that strategies to enhance OHC survival will be necessary to complement Ntf3 as a therapeutic for ARHL. Importantly, despite the threshold shifts observed in the aged Ntf3-overexpressing mice, there was significant improvement in the amplitudes of all peaks of the ABR, suggesting that cochlear Ntf3 supplementation by itself can improve auditory neural function during aging, even when facing significant OHC loss or dysfunction.

Whereas good cochlear thresholds are key to maintaining the audibility of sounds, healthy neural populations are key to maintaining the intelligibility of sounds, especially in difficult listening situations, where the central nervous system must pool responses from large population of auditory neurons to improve the signal-to-noise ratio. Histopathological studies of human autopsy specimens have demonstrated a dramatic loss of auditory nerve fibers in “normal-aging” adults, despite the survival of most of their IHC targets (Wu et al., 2019). It is likely that this primary neural degeneration is a key contributor to the classic complaint of those with ARHL, i.e. problems understanding speech in a noisy environment. The present results suggest that neurotrophin therapies could prevent or slow much of this age-related degradation in hearing abilities.

We previously showed that IHC-SGN synapse density and ABR peak 1 amplitude strongly correlate in young mice (Wan et al., 2014). Here, we demonstrate that these variables also correlate with Ntf3 expression levels in aged mice. Our results also show that activating Ntf3 overexpression in the middle-aged cochlea rapidly increases the amplitude of cochlear sound-evoked potentials without altering ABR or DPOAE thresholds. Remarkably, this does not occur when Ntf3 is overexpressed in young adults (2-month-old). This difference likely reflect the age-related decline in endogenous Ntf3 expression (Fig. 4 and (Sugawara et al., 2007)), i.e., Ntf3 overexpression enhances IHC-SGN synapses acutely only after the endogenous Ntf3 expression is reduced due to aging. Together, with the finding that Ntf3 overexpression in middle age can prevent subsequent loss of IHC-SGN synapses, our results suggest that the age-related decline in endogenous Ntf3 expression contributes to age-related cochlear synaptopathy and reductions in ABR peak 1 amplitudes.

Our present findings provide proof-of-concept that increasing cochlear Ntf3 levels in the middle-aged ear can prevent the age-related degradation of auditory responses, maintaining IHC-SGN synapses and their function. Our previous studies on noise-induced synaptopathy showed that increasing cochlear Ntf3 levels promotes synapse preservation or regeneration when the neurotrophin is provided via transgenesis (Wan et al., 2014), direct Ntf3 application to the round window (Suzuki, Corfas, & Liberman, 2016) or AAV-mediated gene transfer (Hashimoto et al., 2019). Thus, we anticipate that both protein delivery and gene therapy will work also for ARHL, and new approaches for Ntf3 delivery to the round window have been developed (Gunewardene et al., 2022; J. Wang et al., 2021). Of course, a virus-mediated gene therapy approach would have the advantage of long-lasting expression that is likely necessary to preserve IHC-SGN synapses over years, similar to the effects we observed here in transgenic mice. However, other studies indicate that careful control of virus titer and levels of Ntf3 overexpression will be necessary to avoid negative side effects. For example, we found that while relatively modest levels of Ntf3 overexpression (2-4-fold over control) via AAV-mediated gene therapy (Hashimoto et al., 2019) or transgenesis (Wan et al., 2014) promote recovery of IHC-SGN synapses after noise exposure in young mice, higher Ntf3 overexpression levels (50 -100 fold over control) actually increases noise-induced synaptopathy (Hashimoto et al., 2019).

Synaptopathy is not only involved in the pathogenesis of ARHL but has been proposed to play early and critical roles in a number of neurological and neurodegerative disorders (Henstridge, Pickett, & Spires-Jones, 2016). Thus, our findings in the cochlea raise the possibility that increasing the availability of factors that regulate synaptogenesis in the central nervous system during development might serve as therapies for degenerative disorders by slowing or preventing age-related synaptopathy.

## Experimental Procedures

### Animals and tamoxifen treatment

To overexpress Ntf3 from adult supporting cells, we crossed *Slc1a3/CreER*^*T*^ mice (*Slc1a3 CreER*^*T*^; JAX stock no. 012586) with *Ntf3*^*stop*^ mice. The conditional Ntf3 overexpression transgene is under regulation of the synthetic CAGGS promoter/enhancer/intron followed by a loxP-STOP-loxP cassette. *Slc1a3/CreER*^*T*^ mice (Y. Wang et al., 2012) were obtained from Jackson Laboratory while the engineering of the *Ntf3*^*stop*^ mouse was described elsewhere (Wan et al., 2014). *Ntf3*^*stop*^*:Slc1a3/CreER*^*T*^ mice and their controls were on a mixed background of C57BL/6 and FVB/N, and were genotyped to ensure that they do not carry the *ahl* cadherin 23 allele associated with early onset progressive hearing loss present in C57BL/6 mice (Noben-Trauth, Zheng, & Johnson, 2003). Cre-negative littermates (*Ntf3*^*stop*^) were used as control group. For the study, either 8-week-old or 60-week-old mice were gavaged with tamoxifen (200 mg/kg/day) for 3 days. All animal procedures were approved by Institutional Animal Care and Use Committee of the University of Michigan, and all experiments were performed in accordance with relevant guidelines and regulations.

### Physiological analyses

Inner ear physiology, including auditory brainstem responses (ABRs, the summed activity of auditory afferent pathways to short tone bursts), and distortion product otoacoustic emissions (DPOAEs), was performed on mice anesthetized with a mixture of ketamine (100 mg/kg, i.p.) and xylazine (20 mg/kg, i.p.). For the middle-aged mice the first recording was performed at 59 weeks of age (baseline), followed by tamoxifen treatment at 60 weeks of age and additional measurements at 61, 62, 64 and 95 weeks of age. For the young adult mice the first recording was performed at 7 weeks of age (baseline), followed by tamoxifen treatment at 8 weeks of age and additional measurement at 9 weeks of age. For ABR recordings, 3 needle electrodes were placed into the skin at the dorsal midline: one close to the neural crest, one behind the left pinna, and one at the base of the tail (ground). ABR potentials were evoked with 5 ms tone pips (0.5 ms rise-fall, with a cos^2^ envelope, at 40/s) delivered to the eardrum at log-spaced frequencies from 5.6 to 42.25 kHz. The response was amplified (10,000X) and filtered (0.3–3 kHz) with an analog-to-digital board in a PC-based data-acquisition system. Sound pressure level was raised in 5 dB steps from 20 to 80 dB SPL. At each level, 1024 responses were averaged (with stimulus polarity alternated) after ‘artifact rejection’ above 15 µV. The DPOAEs, in response to two primary tones of frequencies f1 and f2, were recorded at (2 × f1) − f2, with f2/f1 = 1.2, and the f2 level 10 dB lower than the f1 level. Stimuli were raised in 5 dB steps from 20 to 80 dB. The ear-canal sound pressure was amplified and digitally sampled at 4 µs intervals. DPOAE thresholds were defined as the lower SPL where (2f1-f2) - (2f1-f2Nse) > 0. These acoustic signals, generated by outer hair cells and measurable in the ear canal, are useful for differential diagnosis: attenuation of ABRs without a change in DPOAEs provides strong evidence for cochlear synaptic or neural dysfunction (Kujawa & Liberman, 2009). Both ABR and DPOAE recordings were performed using the EPL Cochlear Function Test Suite (Mass Eye and Ear, Boston, MA, USA). ABR thresholds, ABR peak 1 amplitudes and latencies, ABR waveforms and DPOAE thresholds were analyzed with ABR peak Analysis software (Mass Eye and Ear, Boston, MA, USA) and Microsoft Excel.

### RNA isolation and quantitative RT-PCR

Mice (10- and 95-weeks-old) were euthanized by cervical dislocation followed by decapitation and the inner ear temporal bones were harvested. Total RNA was purified from whole individual inner ears using RNA extraction kit and Qiazol Reagent (RNeasy mini kit; Qiagen, Germany). DNase treatment was performed immediately after total RNA purification (RNase-free; Qiagen). First strand cDNA was prepared from 400 ng of total RNA and the complementary DNA strand was synthesized using iScript cDNA synthesis kit (Bio-Rad, #1708891, USA), according to the manufacturers protocol. Quantitative RT-PCR was carried out by triplicate using cDNA aliquots derived from each of 9 inner ears (10-week-old *Ntf3*^*stop*^), 7 inner ears (95-week-old *Ntf3*^*stop*^) and 11 inner ears (95-week-old *Ntf3*^*stop*^*:Slc1a3/CreER*^*T*^). The 10 μl reaction contained 5 μl of SYBR Green supermix, 6 pmol of each forward and reverse primer (0.6 μl), 1.9 μl nuclease-free of water and 2.5 μl of cDNA sample. cDNA samples were amplified using iTaq Universal SYBR® Green supermix (Bio-Rad, # 172-5121, USA). Upon PCR completion, PCR products were melted by gradually increasing the temperature in 0.5°C steps. Each sample had a matched “no-RT” control which was tested simultaneously. Water instead of complementary DNA was used as a negative control. RT-qPCR was performed on a CFX-96 Bio-Rad reverse transcription polymerase chain reaction detection system (Hercules, CA, USA). Relative transcript levels of Ntf3 gene were determined by a comparative cycle threshold (Ct) method and relative gene copy number was calculated as normalized gene expression, defined as described previously (Stankovic & Corfas, 2003). Changes in mRNA expression were calculated as relative expression (arbitrary units) respective to the control group. Ribosomal protein L19 (RPL19) was used as the housekeeping gene for normalization. Primers pairs were synthesized by IDT (Coralville, IA, USA): Ntf3, F 5’ GCCCCCTCCCTTATACCTAATG 3’; R: 5’ CATAGCGTTTCCTCCGTGGT 3’; Rpl19, F: 5’ ACCTGGATGAGAAGGATGAG 3’; R: 5’ ACCTTCAGGTACAGGCTGTG 3’.

### Immunostaining for hair cells and synaptic counts

Inner ear tissues from 10 and 95-week-old mice were dissected and fixed in 4% paraformaldehyde in 0.01M phosphate-buffered saline (PBS) for 2 h at room temperature, followed by decalcification in 5% EDTA at 4 °C for 5 days. Cochlear tissues were microdissected and permeabilized by freeze–thawing in 30% sucrose. The microdissected tissues were incubated in blocking buffer containing 5% normal goat serum and 0.3 % Triton X-100 in PBS for 1 h. Tissues were then incubated in primary antibodies (diluted in 1 % normal goat serum and 0.3 % Triton X-100 in PBS) at 37 °C overnight. The primary antibodies used in this study were as follows: anti-Ctbp2 (BD Biosciences, San Jose, CA; 1:200; catalog no. 612044), anti-GluR2 (Millipore, Billerica, MA; 1:1000; catalog no. MAB397) and anti-MyoVIIa (Proteus Biosciences, Ramona, CA; 1:100; catalog no. 25-6790). Tissues were then incubated with appropriate Alexa Fluor-conjugated fluorescent secondary antibodies (Invitrogen, Carlsbad, CA; 1:1000 diluted in 1 % normal goat serum and 0.3 % Triton X-100 in PBS; AF488 IgG2a catalog no. A-21131; AF568 IgG1 catalog no. A-21124; AF647 IgG catalog no. A-21244) for 1 h at room temperature. The tissues were mounted on microscope slides in ProLong Diamond Antifade Mountant (Thermo Fisher Scientific). All pieces of each cochlea were imaged at low power (10X magnification) to convert cochlear locations into frequency using a custom plug-in to ImageJ (1.53c NIH, MD) available at the website of the Eaton-Peabody Laboratories (EPL). Cochlear tissues from 8 to 32 kHz regions were used for further analyses. Confocal z-stacks of cochlear tissues were taken using a Leica SP8 confocal microscope. Images for hair cell counts were taken under 40X magnification. For inner hair cell synapse counts, z-stacks (0.3 µm step size) were taken under 63X (+2.4X optical zoom) magnification spanning the entire IHC height so that all the synaptic specializations were imaged. Imaging and analyses of cochlear hair cells and synapses were performed as previously described in (Wan et al., 2014). Briefly, the number of inner and outer hair cells at specific cochlear regions was determined based on the MyoVIIa channel. Leica Application Suite X software (LASX, Leica microsystem) was used for image processing of z-stacks. All immunofluorescence images shown are representative of at least three individual pictures taken from adjacent areas correspondent to the frequency map and collected from 5-6 mice from each group. ImageJ/Fiji software (version 1.53c, NIH, MD) was used for image processing and three-dimensional reconstruction of z-stacks. CtBP2 and GluR2 puncta in each image stacks were then captured and counted manually using ImageJ/Fiji software multi-point counter tool. Synaptic counts of each z-stack were divided by the number of inner hair cells, which could be visualized by staining of MyoVIIa antibody. Each individual image usually contained 8–10 inner hair cells.

### Statistical analysis and correlations

Graphics and statistical tests were performed using GraphPad Prism version 9.3.1 for Windows (GraphPad Software, San Diego, California USA, www.graphpad.com). Statistical differences in auditory physiology (ABR thresholds, DPOAE thresholds and ABR peak 1 amplitudes) were analyzed using two-way ANOVA, followed by Bonferroni’s multiple comparisons test. Statistical differences for confocal microcopy images used for quantification were analyzed using either two-tailed t-test or one-way ANOVA, followed by Tukey’s multiple comparisons test. Statistical differences for Ntf3 mRNA expression were analyzed using Kruskal-Wallis test, followed by Dunn’s multiple comparisons test. Correlations were evaluated using non-parametric Spearman correlation test. All p < 0.05 were considered as statistically significant.

## Acknowledgement

This work was supported by NIH R01DC004820 (GC), R01DC018500 (GC), T32DC005356 Advanced Research Training in Otolaryngology (NDC), R01 DC0188 (MCL).

## Author contributions

designing research studies (LRC, GC), conducting experiments (LRC, LJ, BCB, NC), acquiring data (LRC, LJ, BCB, NC, ASD, DK), data analysis (LRC, BCB, ASD, GC), writing the manuscript (LRC, LJ, BCB, NC, AD, DK, MCL, GC).

## Data availability

The data that support the findings of this study are available from the corresponding author upon reasonable request.

---

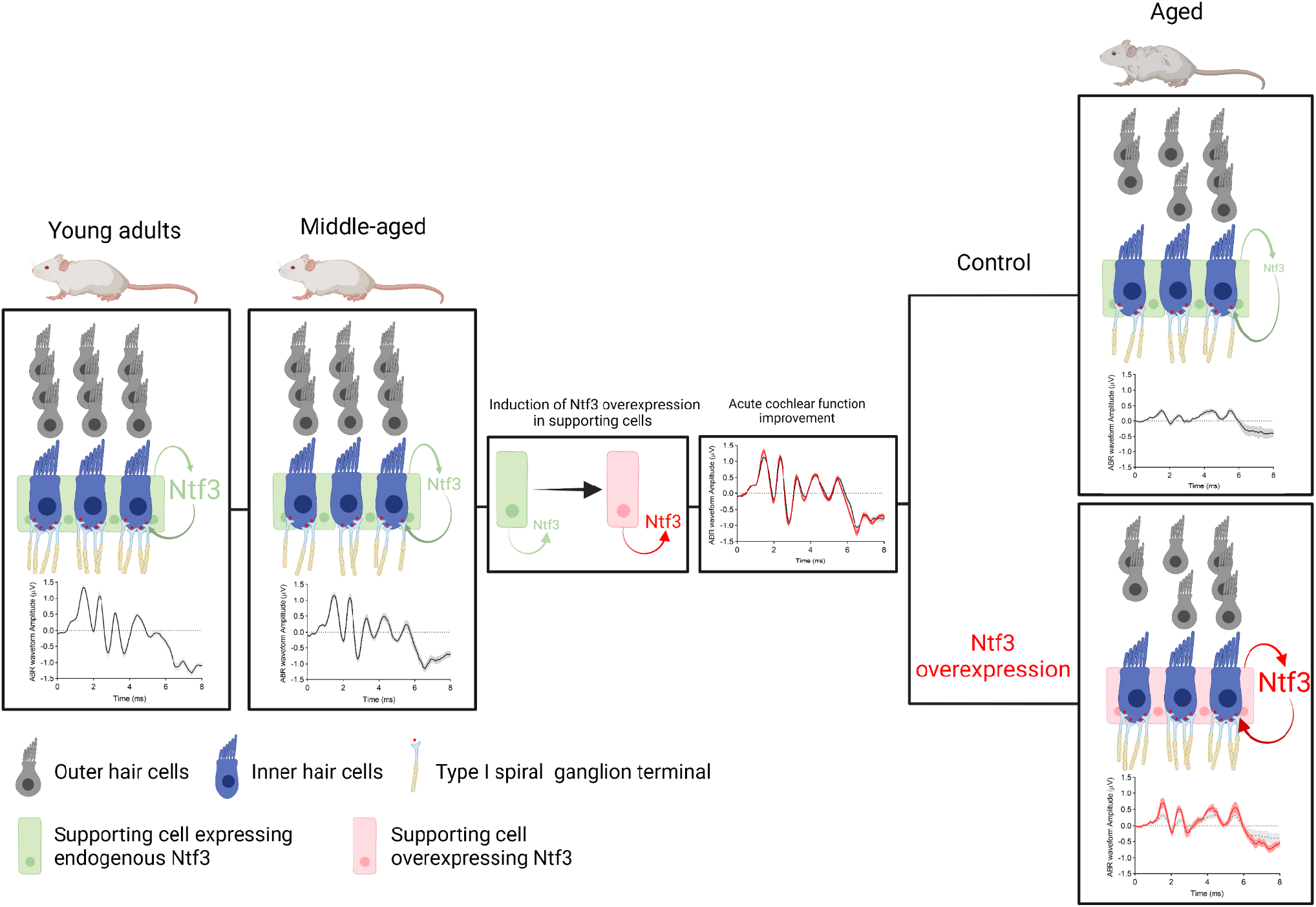

## References

Agrawal, Y., Platz, E. A., & Niparko, J. K. (2008). Prevalence of hearing loss and differences by demographic characteristics among US adults: data from the National Health and Nutrition Examination Survey, 1999-2004. Arch Intern Med, 168(14), 1522–1530. doi:10.1001/archinte.168.14.1522

Bainbridge, K. E., & Wallhagen, M. I. (2014). Hearing loss in an aging American population: extent, impact, and management. Annu Rev Public Health, 35, 139–152. doi:10.1146/annurev-publhealth-032013-182510

Ernfors, P., Van De Water, T., Loring, J., & Jaenisch, R. (1995). Complementary roles of BDNF and NT-3 in vestibular and auditory development. Neuron, 14(6), 1153–1164. doi:10.1016/0896-6273(95)90263-5

Farinas, I., Jones, K. R., Backus, C., Wang, X. Y., & Reichardt, L. F. (1994). Severe sensory and sympathetic deficits in mice lacking neurotrophin-3. Nature, 369(6482), 658–661. doi:10.1038/369658a0

Gunewardene, N., Lam, P., Ma, Y., Caruso, F., Wagstaff, S., Richardson, R. T., & Wise, A. K. (2022). Pharmacokinetics and biodistribution of supraparticle-delivered neurotrophin 3 in the guinea pig cochlea. J Control Release, 342, 295–307. doi:10.1016/j.jconrel.2021.12.037

Hashimoto, K., Hickman, T. T., Suzuki, J., Ji, L., Kohrman, D. C., Corfas, G., & Liberman, M. C. (2019). Protection from noise-induced cochlear synaptopathy by virally mediated overexpression of NT3. Sci Rep, 9(1), 15362. doi:10.1038/s41598-019-51724-6

Henstridge, C. M., Pickett, E., & Spires-Jones, T. L. (2016). Synaptic pathology: A shared mechanism in neurological disease. Ageing Res Rev, 28, 72–84. doi:10.1016/j.arr.2016.04.005

Huddle, M. G., Goman, A. M., Kernizan, F. C., Foley, D. M., Price, C., Frick, K. D., & Lin, F. R. (2017). The Economic Impact of Adult Hearing Loss: A Systematic Review. JAMA Otolaryngol Head Neck Surg, 143(10), 1040–1048. doi:10.1001/jamaoto.2017.1243

Knipper, M., Zimmermann, U., Rohbock, K., Kopschall, I., & Zenner, H. P. (1996). Expression of neurotrophin receptor trkB in rat cochlear hair cells at time of rearrangement of innervation. Cell Tissue Res, 283(3), 339–353. doi:10.1007/s004410050545

Kohrman, D. C., Borges, B. C., Cassinotti, L. R., Ji, L., & Corfas, G. (2021). Axon-glia interactions in the ascending auditory system. Dev Neurobiol, 81(5), 546–567. doi:10.1002/dneu.22813

Kujawa, S. G., & Liberman, M. C. (2006). Acceleration of age-related hearing loss by early noise exposure: evidence of a misspent youth. J Neurosci, 26(7), 2115–2123. doi:10.1523/JNEUROSCI.4985-05.2006

Kujawa, S. G., & Liberman, M. C. (2009). Adding insult to injury: cochlear nerve degeneration after “temporary” noise-induced hearing loss. J Neurosci, 29(45), 14077–14085. doi:10.1523/JNEUROSCI.2845-09.2009

Livingston, G., Huntley, J., Sommerlad, A., Ames, D., Ballard, C., Banerjee, S., … Mukadam, N. (2020). Dementia prevention, intervention, and care: 2020 report of the Lancet Commission. Lancet, 396(10248), 413–446. doi:10.1016/S0140-6736(20)30367-6

Makary, C. A., Shin, J., Kujawa, S. G., Liberman, M. C., & Merchant, S. N. (2011). Age-related primary cochlear neuronal degeneration in human temporal bones. J Assoc Res Otolaryngol, 12(6), 711–717. doi:10.1007/s10162-011-0283-2

Noben-Trauth, K., Zheng, Q. Y., & Johnson, K. R. (2003). Association of cadherin 23 with polygenic inheritance and genetic modification of sensorineural hearing loss. Nat Genet, 35(1), 21–23. doi:10.1038/ng1226

Powell, D. S., Oh, E. S., Lin, F. R., & Deal, J. A. (2021). Hearing Impairment and Cognition in an Aging World. J Assoc Res Otolaryngol, 22(4), 387–403. doi:10.1007/s10162-021-00799-y

Sergeyenko, Y., Lall, K., Liberman, M. C., & Kujawa, S. G. (2013). Age-related cochlear synaptopathy: an early-onset contributor to auditory functional decline. J Neurosci, 33(34), 13686–13694. doi:10.1523/JNEUROSCI.1783-13.2013

Stankovic, K. M., & Corfas, G. (2003). Real-time quantitative RT-PCR for low-abundance transcripts in the inner ear: analysis of neurotrophic factor expression. Hear Res, 185(1-2), 97–108. doi:10.1016/s0378-5955(03)00298-3

Sugawara, M., Murtie, J. C., Stankovic, K. M., Liberman, M. C., & Corfas, G. (2007). Dynamic patterns of neurotrophin 3 expression in the postnatal mouse inner ear. J Comp Neurol, 501(1), 30–37. doi:10.1002/cne.21227

Suzuki, J., Corfas, G., & Liberman, M. C. (2016). Round-window delivery of neurotrophin 3 regenerates cochlear synapses after acoustic overexposure. Sci Rep, 6, 24907. doi:10.1038/srep24907

Viana, L. M., O’Malley, J. T., Burgess, B. J., Jones, D. D., Oliveira, C. A., Santos, F., … Liberman, M. C. (2015). Cochlear neuropathy in human presbycusis: Confocal analysis of hidden hearing loss in post-mortem tissue. Hear Res, 327, 78–88. doi:10.1016/j.heares.2015.04.014

Wan, G., Gomez-Casati, M. E., Gigliello, A. R., Liberman, M. C., & Corfas, G. (2014). Neurotrophin-3 regulates ribbon synapse density in the cochlea and induces synapse regeneration after acoustic trauma. Elife, 3. doi:10.7554/eLife.03564

Wang, J., Youngblood, R., Cassinotti, L., Skoumal, M., Corfas, G., & Shea, L. (2021). An injectable PEG hydrogel controlling neurotrophin-3 release by affinity peptides. J Control Release, 330, 575–586. doi:10.1016/j.jconrel.2020.12.045

Wang, Y., Rattner, A., Zhou, Y., Williams, J., Smallwood, P. M., & Nathans, J. (2012). Norrin/Frizzled4 signaling in retinal vascular development and blood brain barrier plasticity. Cell, 151(6), 1332–1344. doi:10.1016/j.cell.2012.10.042

Wu, P. Z., Liberman, L. D., Bennett, K., de Gruttola, V., O’Malley, J. T., & Liberman, M. C. (2019). Primary Neural Degeneration in the Human Cochlea: Evidence for Hidden Hearing Loss in the Aging Ear. Neuroscience, 407, 8–20. doi:10.1016/j.neuroscience.2018.07.053

Ylikoski, J., Pirvola, U., Moshnyakov, M., Palgi, J., Arumae, U., & Saarma, M. (1993). Expression patterns of neurotrophin and their receptor mRNAs in the rat inner ear. Hear Res, 65(1-2), 69–78. doi:10.1016/0378-5955(93)90202-c

